# Forskolin reverses the O-GlcNAcylation dependent decrease in GABA_A_R current amplitude at hippocampal synapses possibly at a neurosteroid site on GABA_A_Rs

**DOI:** 10.1101/2024.03.06.583612

**Authors:** Shekinah Phillips, John C. Chatham, Lori L. McMahon

## Abstract

GABAergic transmission is influenced by post-translational modifications, like phosphorylation, impacting channel conductance, allosteric modulator sensitivity, and membrane trafficking. O-GlcNAcylation is a post-translational modification involving the O-linked attachment of β–N-acetylglucosamine on serine/threonine residues. Previously we reported an acute increase in O-GlcNAcylation elicits a long-term depression of evoked GABA_A_R inhibitory post synaptic currents (eIPSCs) onto hippocampal principal cells. Importantly, O-GlcNAcylation and phosphorylation can co-occur or compete for the same residue; whether they interact in modulating GABAergic IPSCs is unknown. We tested this by recording IPSCs from hippocampal principal cells and pharmacologically increased O-GlcNAcylation, before or after increasing serine phosphorylation using the adenylate cyclase activator, forskolin. Although forskolin had no significant effect on baseline eIPSC amplitude, we found that a prior increase in O-GlcNAcylation unmasks a forskolin-dependent increase in eIPSC amplitude, reversing the O-GlcNAc-induced eIPSC depression. Inhibition of adenylate cyclase or protein kinase A did not prevent the potentiating effect of forskolin, indicating serine phosphorylation is not the mechanism. Surprisingly, increasing O-GlcNAcylation also unmasked a potentiating effect of the neurosteroids 5α-pregnane-3α,21-diol-20-one (THDOC) and progesterone on eIPSC amplitude, mimicking forskolin. Our findings show under conditions of heightened O-GlcNAcylation, the neurosteroid site on synaptic GABA_A_Rs is accessible to agonists, permitting strengthening of synaptic inhibition.

## Introduction

Gamma-aminobutyric acid type A receptors (GABA_A_Rs) are heteropentameric ligand-gated chloride channels composed of α, β, γ, and sometimes δ subunits that mediate both fast synaptic and tonic inhibition, depending on their synaptic vs extrasynaptic location, respectively^1^. Mutations in specific subunits are linked to epilepsy syndromes ^2^, and gene polymorphisms in specific GABA_A_R subunits associate with neuropsychiatric disorders, including alcohol use disorder ^3^, anxiety ^4^, schizophrenia ^5^, bipolar disorder ^5^, and even major depressive disorder ^6^, including postpartum depression ^7^.

For decades it has been appreciated that neurosteroids mediate their sedative hypnotic and anxiolytic effects via positive allosteric binding to specific GABA_A_R subunits, particularly extrasynaptic GABA_A_Rs containing α5, α4 or δ subunits ^8,9^. GABA_A_R function is also potently modulated by serine phosphorylation ^10^. For example, protein kinase A (PKA) mediated phosphorylation of specific serines on synaptic GABA_A_Rs induced by application of the adenylate cyclase activator, forskolin, bidirectionally modulates GABA-gated current amplitude and induces endocytosis depending on the neuron type and phosphorylated residue^8,11–14^. Importantly, the phosphorylation state has direct consequences on potency of allosteric modulation of GABA_A_Rs by neurosteriods, barbiturates, and benzodiazepines in a subunit-specific manner ^10^.

The post-translational modification of proteins by β-N-acetylglucosamine via an O-linkage on serine and threonine residues (O-GlcNAcylation), can modulate protein phosphorylation by competing directly with phosphorylation for the same residues, or indirectly via modification of other sites thereby changing protein structure and protein-protein interactions. In addition, many kinases are modified by O-GlcNAc and this can regulate their function ^15^. Both O-GlcNAcylation and phosphorylation are dynamic, reversible, and ubiquitous. While many kinases and phosphatases exist, O-GlcNAc is tightly regulated by a single enzyme pair, OGT (O-GlcNAc transferase) and OGA (O-GlcNAcase), which add and remove O-GlcNAc from serine/threonine residues, respectively. Of note, these enzymes are highly expressed in hippocampus ^16,17^. The OGT substrate, UDP-GlcNAc, is generated by the hexosamine biosynthetic pathway (HBP) via glucose metabolism potentially linking this modification to nutrient availability ^18,19^.

Our lab recently reported that O-GlcNAcylation can be increased within minutes by exposing hippocampal brain slices to the HBP substrate glucosamine (GlcN), the OGA inhibitor thiamet-G (TMG) or in combination and this leads to depression of excitatory synaptic transmission ^20,21^. More recently, we reported that pharmacologically increasing O-GlcNAc using GlcN and TMG in combination or GlcN alone induces a long-lasting depression of GABA_A_R-mediated inhibitory postsynaptic currents (IPSCs) at hippocampal synapses within minutes, and decreases the amplitude and frequency of spontaneous IPSCs (sIPSCs) ^22^. In *Oga +/−* mice, where O-GlcNAc is chronically elevated, inhibitory synaptic transmission was reduced in medial prefrontal cortex, and this was rescued by OGA overexpression via adeno-associated viral injection ^23^. Additionally, the *Oga +/−* mice exhibited an antidepressant-like behavior, which was also reversed by viral OGA overexpression. Collectively, acute and chronic elevation in O-GlcNAcylation *in vitro* and *in vivo* depresses GABA_A_R-mediated synaptic inhibition. These findings highlight O-GlcNAcylation as a critical regulator of the efficacy of synaptic neuronal inhibition, in both physiological and pathophysiological conditions of elevated O-GlcNAcylation.

Although serine phosphorylation ^10,11^ and O-GlcNAcylation ^22^ are fundamental modulators of the strength of GABA_A_R-mediated synaptic inhibition, no study has examined whether these modifications interact in modifying the efficacy of synaptic inhibition. We investigated this possibility using electrophysiology in hippocampal slices and pharmacologically increased O-GlcNAc using glucosamine (GlcN) and thiamet-G (TMG) in combination. We also increased serine phosphorylation using the adenylate cyclase activator forskolin. Unexpectedly, we found that a prior increase in O-GlcNAcylation, which induces depression of GABA_A_R-mediated eIPSC amplitude, unmasks a forskolin-dependent increase in eIPSC amplitude, even though forskolin had no effects when applied in the absence of increased O-GlcNAcylation. Surprisingly, inhibition of adenylate cyclase or PKA did not prevent the potentiating effect of forskolin on eIPSC amplitude, indicating that serine phosphorylation is not the mechanism. Similar to findings in a study in carp amacrine-like cells showing that forskolin binds to a GABA_A_R neurosteroid site^24^, we found that the neurosteriods, THDOC and progesterone potentiate the IPSC amplitude following a prior increase in O-GlcNAcylation, mimicking the effect of forskolin. These findings suggest that O-GlcNAcylation promotes neurosteroid site accessibility on GABA_A_Rs thereby reversing the depressive effect of O-GlcNAcylation and strengthening synaptic inhibition.

## Results

### An acute increase in O-GlcNAcylation reduces inhibitory post-synaptic currents on CA1 pyramidal cells

Using hippocampal slices and Western blot analysis, we previously reported that an acute 10 min exposure to the HBP substrate GlcN (100 µM or 5 mM) or to the OGA inhibitor TMG (1 µM) induces a significant, global increase in protein O-GlcNAcylation (O-GlcNAc) ^20,21^. This same 10 min exposure induces a long-term depression (LTD) at both excitatory CA3-CA1 synapses ^20,21^, and inhibitory synapses^22^ during electrophysiological recordings. Additionally, increasing protein O-GlcNAc reduces the amplitude and frequency of spontaneous IPSC (sIPSCs) recorded from CA1 pyramidal cells and dentate granule cells (DGCs). Here, we confirm that a 10 min bath application of GlcN (5mM) and TMG (1µM) (GlcN+TMG) to pharmacologically increase O-GlcNAcylation during whole-cell voltage clamp recordings from CA1 pyramidal cells significantly depresses the sIPSC amplitude (Figure 1Aii, cumulative probability distribution, p < 0.0001, KS D value =0.17, Kolmogorov-Smirnov test; inset: p < 0.0001, Wilcoxon matched-pairs signed rank test) and increases the interevent interval, reflecting a decrease in sIPSC frequency (Figure 1Aiii, cumulative probability distribution, p < 0.0001, KS D value = 0.23, Kolmogorov-Smirnov test; inset: p < 0.0001, Wilcoxon matched-pairs signed rank test). Our previous work also showed that the amplitude of miniature IPSCs (mIPSCs) was significantly decreased by increasing O-GlcNAcylation^22^, suggesting this synaptic depression is through a postsynaptic mechanism.

**Figure 1.**
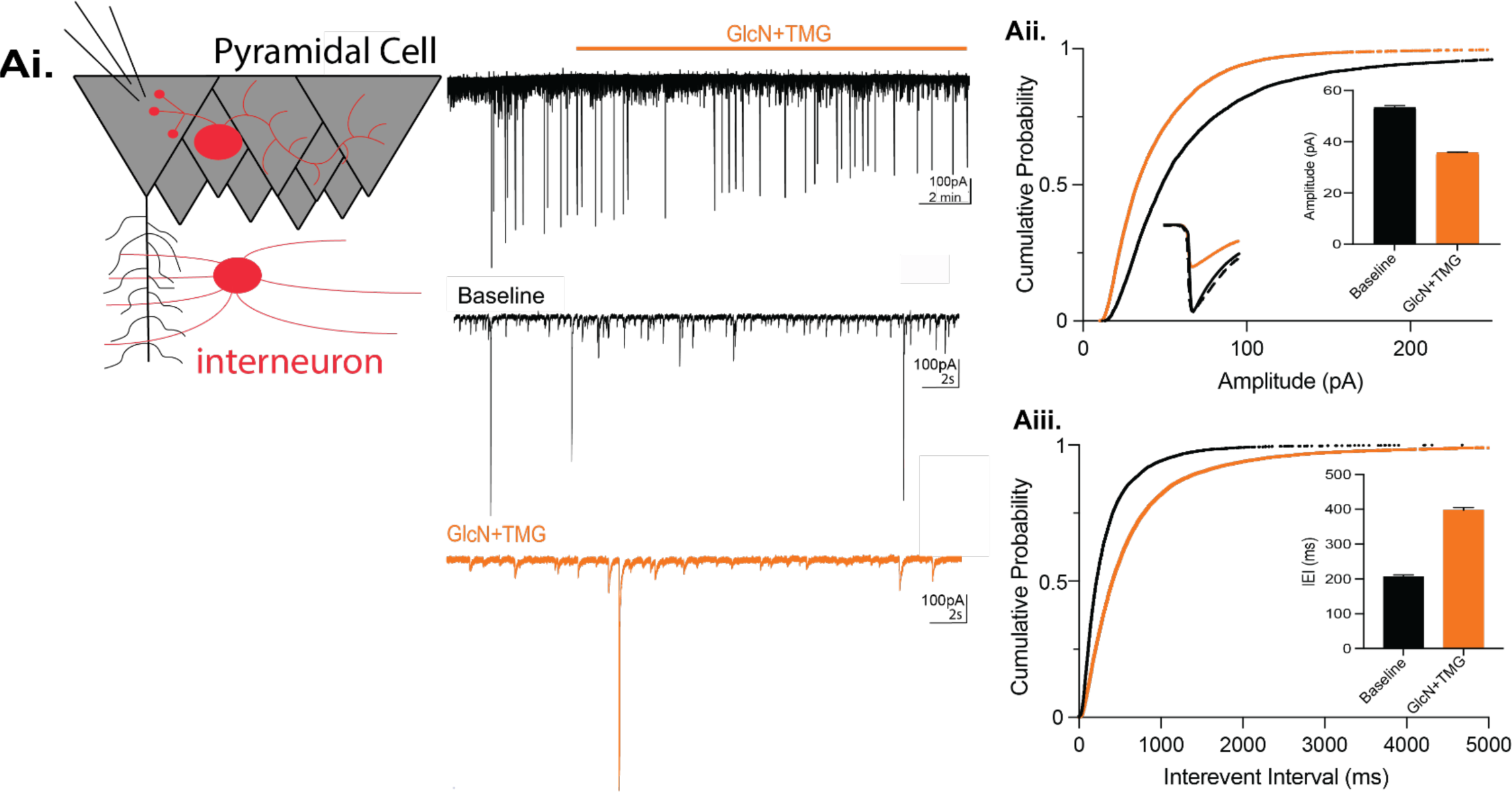
Acute increase in O-GlcNAcylation reduces spontaneous IPSCs in hippocampal CA1 pyramidal cells. **Ai.** (left) Schematic illustrating whole-cell recording of CA1 pyramidal cells (right) representative sIPSC trace showing baseline and GlcN + TMG application (top)and expanded time scale (bottom) during baseline (black) and GlcN + TMG application (orange). **Aii.** Cumulative probability distribution of sIPSC amplitude (p < 0.0001, KS D value = 0.17, Kolmogorov-Smirnov test); *inset*: (left) average sIPSC trace before (black) and after (orange) GlcN + TMG;scaled trace (dashed) shows no change in rise or decay time of depressed sIPSC. *inset*: (right) bar chart showing average (±SEM) sIPSC amplitude. Baseline: 69.7 ± 0.8 pA, GlcN + TMG: 43.5 ± 0.5 pA (p < 0.0001, Wilcoxon matched-pairs signed rank test, n = 9 cells, 6 rats). **Aiii.** Cumulative probability distribution of sIPSC interevent interval (p < 0.0001, KS D value = 0.225, Kolmogorov-Smirnov test). Baseline: 344.5 ± 4.7 ms, GlcN + TMG: 767.3 ± 16.6 ms, *inset*: (p < 0.0001, Wilcoxon matched-pairs signed rank test, n = 9 cells, 6 rats).

### The O-GlcNAc-induced depression of synaptic inhibition is not prevented by an actin stabilizer, but is partially dependent upon dynamin-mediated GABA_A_R endocytosis

Because serine phosphorylation can cause GABA_A_R endocytosis in a subunit and serine specific manor ^10,25,26^ by analogy, we speculated that the O-GlcNAc-induced LTD of synaptic inhibition (or O-GlcNAc iLTD) we previously reported^22^ is occurring via GABA_A_R endocytosis. This possibility is supported by findings from our lab and others that O-GlcNAcylation of the GluA2 AMPAR subunit leads to long-term depression of excitatory transmission (O-GlcNAc LTD) at hippocampal CA3-CA1 synapses^20^ and causes AMPAR endocytosis ^27^.

Therefore, to determine if increasing O-GlcNAc induces GABA_A_R endocytosis during expression of O-GlcNAc iLTD, two experiments were performed. First, we tested whether interfering with actin prevents O-GlcNAc iLTD. We recorded CA1 pyramidal cells (Cs Gluconate pipette solution; E_Cl−_ = −60 mV) and included the actin stabilizer, jasplakinolide (jasp) ^28^, in the pipette solution during whole-cell voltage clamp recordings. We interleaved experiments using pipette solution without jasplakinolide to ensure successful expression of O-GlcNAc iLTD. After a 15-min baseline recording, GlcN+TMG was applied for 10 min to induce O-GlcNAc iLTD. With or without jasplakinolide, successful iLTD was induced (Fig. 2Ai, O-GlcNAc iLTD – jasp: 69.2 ± 5.4% of baseline transmission, p=0.002, paired t-test; Fig. 2Aii, O-GlcNAc iLTD + jasp: 75.2 ± 3.4% of baseline transmission, p=0.0002, paired t-test). However, no significant difference was found between groups (Fig. 2Aiii, p=0.59, One-way ANOVA), indicating that the O-GlcNAc iLTD is not caused by actin mediated GABA_A_R endocytosis.

**Figure 2.**
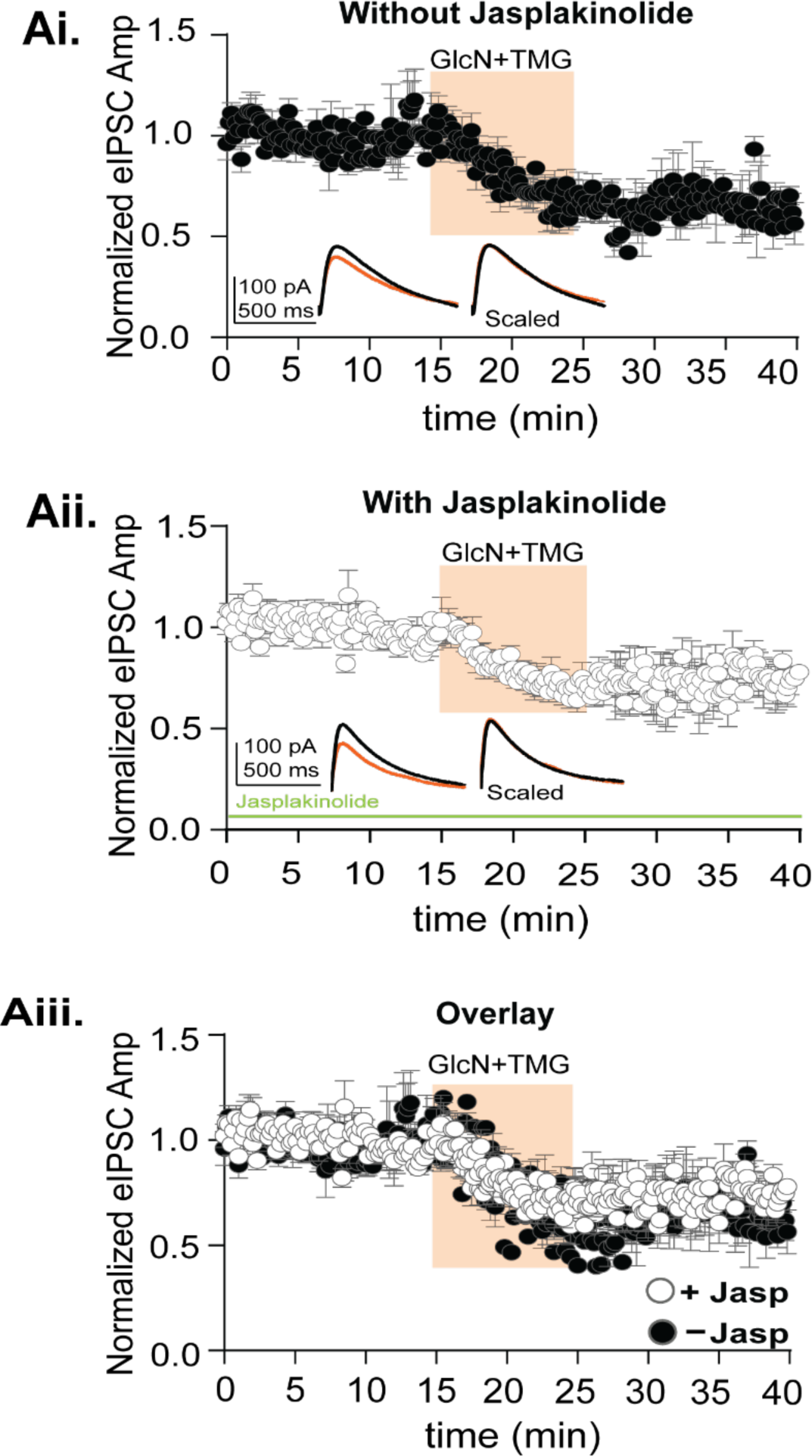
Blocking actin mediated GABA_A_R endocytosis does not prevent O-GlcNAciLTD in CA1 pyramidal cells. **Ai.** Normalized data showing average eIPSC amplitude over time exposed to GlcN+TMG (10 min, orange) after a 15-minute baseline without jasp, (p= 0.005, paired t-test, n = 6 cells, 6 rats). *Inset*: representative eIPSC traces before (black) and after GlcN + TMG (orange) application and scaled average without jasp shows no change in rise or decay time of the eIPSC. **Aii.** Normalized data showing average eIPSC amplitude over time exposed to GlcN+TMG (10 min, orange) after a 15-minute baseline with jasplakinolide included in the pipette solution (jasp) (p= 0.0005, paired t-test, n = 8 cells, 8 rats). *Inset*: representative eIPSC traces before (black) and after GlcN + TMG (orange) application and scaled average with jasp shows no change in rise time or decay of the eIPSC.. **Aiii.** Overlay of the experimental groups showing no significant difference in the average (±SEM) eIPSC amplitude over time (p<0.0001, One-way ANOVA). Gray horizontal bars represent the mean ± SEM.

Next, we tested whether expression of O-GlcNAc iLTD requires dynamin-dependent GABA_A_R endocytosis. To accomplish this, we incubated slices in dynasore (80 μM, 30 min) or DMSO (vehicle) and performed whole-cell recordings from CA1 pyramidal cells in an interleaved fashion. After a 5 min baseline, GlcN+TMG was applied for 10 min to induce iLTD. With or without dynasore, successful iLTD was induced (Fig. 3Ai, O-GlcNAc iLTD – dynasore: 79.3 ± 4.5% of baseline transmission, p= 0.005, paired t-test; Fig. 3Aii, O-GlcNAc iLTD + dynasore: 81.1 ± 13.2% of baseline transmission, p=0.004, paired t-test). However, there were no significant differences between groups (Fig.3Aiii, p=0.08, One-Way ANOVA). Importantly, it was noted that the dataset with dynasore had greater variability during iLTD expression as indicated by the larger error bars between 20-25 mins (Fig. 3Aii). Upon further inspection of individual experiments, we recognized that in some cells (n=4/8), the eIPSC amplitude was potentiated following GlcN+TMG, which would be expected if endocytosis is prevented. Therefore, when the dynasore dataset was separated into those with potentiation and those without, a clear population of cells was revealed that exhibited significant potentiation of the eIPSC amplitude (Fig. 3Aiv, 136.3 ± 10.2% of normalized eIPSC amplitudes following GlcN+TMG, application n= 4 cells, p=0.039, paired t-test), while the remaining population exhibited no change in eIPSC amplitude from the previously depressed level following GlcN+TMG application (Fig. 3Aiv, 73.9 ± 13.5% of normalized eIPSC amplitudes following GlcN+TMG, application n= 4 cells, p=0.13, paired t-test) and from the O-GlcNAC iLTD without dynasore (Fig3Ai) (p=0.131, unpaired t-test). Additionally, there was a significant difference between the potentiated versus the non-potentiated population of normalized eIPSC amplitudes following GlcN+TMG (p=0.03, paired t-test). This result is consistent with the interpretation that in some cells increasing O-GlcNAc induced a dynamin-dependent endocytosis of GABA_A_Rs that could underlie the synaptic depression and in others a different mechanism exists. Furthermore, these findings reaffirm the heterogeneity in GABA_A_Rs that exist at synapses in hippocampus. Additional experiments are needed to fully understand how O-GlcNAc impacts GABA_A_R trafficking.

**Figure 3.**
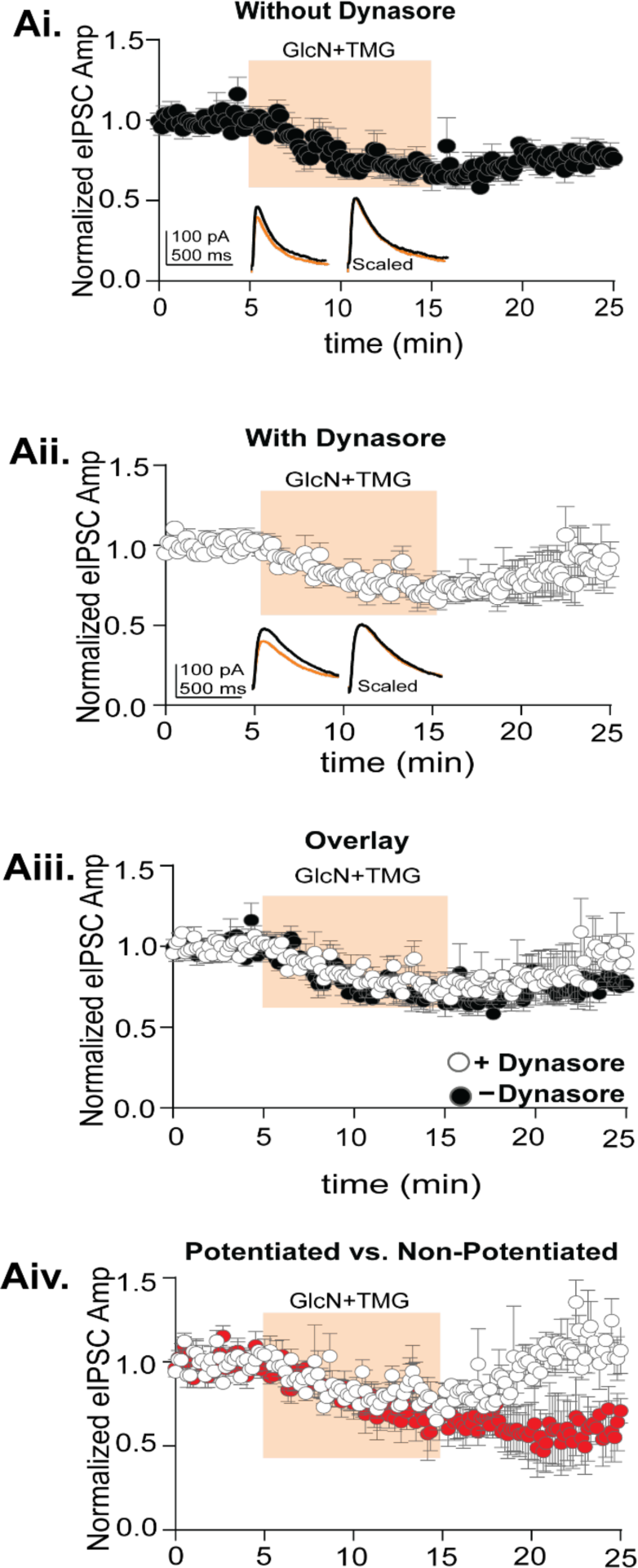
O-GlcNAc iLTD is partially dependent upon dynamin-mediated GABA_A_R endocytosis. **Ai.** Normalized data showing average eIPSC amplitude over time exposed to GlcN+TMG (10 min, orange) after a 15-minute baseline without dynasore (DMSO-vehicle) (p= 0.005, paired t-test, n = 8 cells, 7 rats). *Inset:* representative eIPSC traces before (black) and after GlcN + TMG (orange) application and scaled average shows no difference in rise or decay time of the eIPSC. **Aii.** Normalized data showing average eIPSC amplitude over time exposed to GlcN+TMG (10 min, orange) after a 5-minute baseline in slices incubated in dynasore for 30 min (p= 0.004, paired-test, n = 8 cells, 7 rats). *Inset*: representative eIPSC traces before (black) and after GlcN + TMG (orange) application and scaled average with dynasore. **Aiii.** Overlay of experimental groups showing no significant difference in the average (±SEM) eIPSC amplitude over time (p=0.08, One-way ANOVA). **Aiii.** Separation of two populations in the presence of dynasore where the eIPSC amplitude was significantly increased in one population (white, p=0.0392, paired t-test, n= 4 cells, 4 rats) and with no effect in the other population (red, p=0.13, paired t-test, n= 4 cells, 4 rats). Gray horizontal bars represent the mean ± SEM.

### Possible interactions between phosphorylation and O-GlcNAcylation

Next, we wanted to determine if serine phosphorylation and O-GlcNAcylation interact to affect GABA_A_R-mediated synaptic inhibition and whether an order effect exists. For decades, forskolin has been used to activate adenylate cyclase to drive protein kinase A (PKA) dependent phosphorylation of AMPARs at excitatory synapses in hippocampus, leading to synaptic potentiation ^29–31^. PKA-dependent serine phosphorylation also modulates synaptic inhibition, but the effect is variable depending on the GABA_A_R subunit confirmation ^10^. Therefore, to test whether serine phosphorylation has an impact on subsequent induction of O-GlcNAc iLTD, we recorded from CA1 pyramidal cells (Cs Gluconate pipette solution; E_Cl−_ = −60 mV), and bath applied forskolin (50µM) for 10 min to drive activation of adenylate cyclase and PKA followed by 10 min application of GlcN+TMG to induce O-GlcNAc iLTD. The eIPSC amplitudes during forskolin and GlcN+TMG were normalized to baseline and statistically compared by averaging 30 events during (a) baseline, (b) forskolin and (c) GlcN+TMG using repeated measures (RM) RM-ANOVA and Šídák’s multiple comparisons post hoc test (Fig. 4Ai-iii, p=0.0001, RM ANOVA). We found no significant effect of forskolin compared to baseline transmission (Fig. 4Aii, 89.2 ± 6.9% of baseline transmission, p=0.39), and subsequent application of GlcN+TMG induced significant iLTD (Fig. 4Aii, 64.2 ± 4.4% of baseline transmission, p<0.0001). Despite no significant effect of forskolin on the eIPSC amplitude in the averaged dataset compared to baseline, we want to ensure there was no effect on the magnitude of subsequently induced O-GlcNAc iLTD. Therefore, we normalized the eIPSC amplitudes at the end of the 10 min forskolin application, thereby establishing new baseline from which to measure the O-GlcNAc iLTD magnitude. We found that from this new baseline, subsequent application of GlcN+TMG induced a significant iLTD (74.9 ± 3.8% of new baseline transmission (b-c comparison), p<0.0001, paired t-test) that is not different from the iLTD magnitude under control conditions obtained in Fig. 2Ai in the absence of jasp (Fig. 2Ai, 67.6 ± 5.6% of baseline transmission versus 73.5 ± 4.2% of new baseline transmission p= 0.53, unpaired t-test). In reviewing the data, it is important to note that there was high cell-to-cell variability in eIPSC amplitude during forskolin application, with some cells displaying potentiation and some depression of the eIPSC amplitude, as can be seen by inspection of the individual data points in the bar chart in Fig. 4Aii.

**Figure 4.**
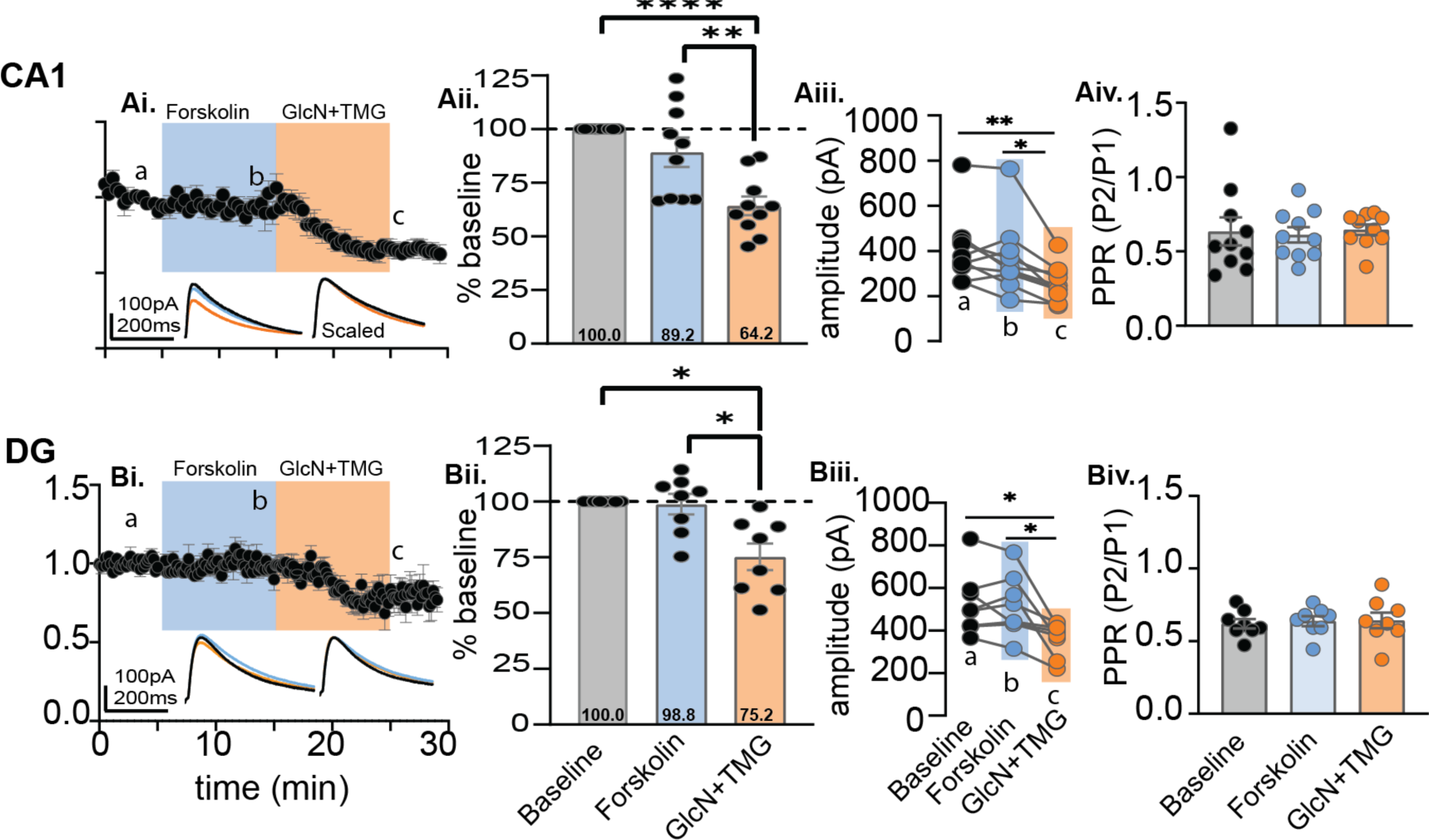
Magnitude of the O-GlcNAc iLTD is unaffected by prior application of forskolin. **Ai.** CA1: Normalized data showing average eIPSC amplitude over time during baseline (5 min), bath application of forskolin (10 min, blue) and GlcN + TMG (10 min, orange). *Inset:* overlaid averaged representative eIPSC traces during baseline (black), forskolin (blue), and GlcN + TMG (orange) (left) and scaled (right) **Aii.** Bar chart illustrates the average affect of forskolin and GlcN+TMG on eIPSC amplitude from the original baseline (dotted line): 89.2% ± 6.9%, p=0.39 (forskolin), 64.2% ± 4.4%, p<0.0001 (GlcN+TMG) and each other (p=0.0023); (p=0.0001, RM-ANOVA). **Aiii.** For each individual experiment in Ai, line graphs show the average eIPSC amplitude during the last the 5 min in each condition corresponding to a,b,c in Ai (p=0.0002, RM-ANOVA, n = 10 cells, 9 rats), Šídák’s: a-c (p=0.002) and b-c (p=0.01). Baseline: 404.9 ± 47.9 pA, forskolin: 366.5 ± 50.2 pA, GlcN+ TMG: 256.6 ± 25.0 pA (mean ± SEM). **Aiv.** Paired-Pulse Ratio (100 ms inter-pulse interval) between baseline, forskolin and GlcN+TMG showed no significant difference (p=0.69, RM-ANOVA). **Bi.** Dentate: Normalized data showing average eIPSC amplitude over time during baseline, bath application of forskolin (blue), and GlcN+TMG (orange). *Inset*: overlaid averaged representative eIPSC traces during baseline (black) forskolin (blue), GlcN + TMG (orange) (left) and scaled (right). **Bii.** Bar chart illustrates the average affect of forskolin and GlcN+TMG from the original baseline (dotted line): 98.8% ± 4.6%, p=0.98 (forskolin) and 75.2% ± 6.0%, p=0.0128 (GlcN+TMG) and each other (p=0.0453); (p=0.004, RM-ANOVA). **Biii.** For each individual experiment in Bi, line graphs show the average amplitudes for last 5 min in each condition corresponding to a,b,c in Bi (p = 0.005, RM-ANOVA, n = 8 cells, 8 rats), Šídák’s: a-c (p=0.029) and b-c (p=0.040). Baseline: 523.7 ± 51.7 pA, forskolin: 514.9 ± 50.5 pA, GlcN+ TMG: 360.2 ± 28.3 pA (mean ± SEM). **Biv.** Paired-Pulse Ratio between baseline, forskolin and GlcN+TMG showed no significant difference (p=0.46, RM-ANOVA). Gray horizontal bars represent the mean ± SEM. Šídák’s post hoc depicted with asterisks.

To determine if this finding in CA1 is generalizable, we performed the same experiment in dentate granule cells (DGCs). Similar to CA1, we recorded from DGCs (Cs Gluconate pipette solution; E_Cl−_ = −60 mV), and bath applied forskolin (50µM) for 10 min followed by 10 min of GlcN+TMG to induce O-GlcNAc iLTD. Again, the eIPSC amplitudes during forskolin and GlcN+TMG were normalized to baseline and statistically compared by averaging 30 events during (a) baseline, (b) forskolin and (c) GlcN+TMG (Fig. 4Bi, p=0.0041, RM-ANOVA). Using Šídák’s multiple comparisons post hoc test, we found no significant change in eIPSC amplitude during forskolin application compared to baseline (Fig. 4Bi, 98.8 ± 4.6% of baseline transmission, p = 0.98), with subsequent GlcN+TMG application inducing significant iLTD (Fig. 4Bii, 75.2 ± 6.0% of baseline, p=0.013). In addition, there was a significant difference between baseline vs. GlcN+ TMG (Fig. 4Biii, p=0.029) and between forskolin vs GlcN+TMG (Fig. 4Biii, p=0.040). Also, as before, to measure the magnitude of the O-GlcNAc iLTD after forskolin, we re-normalized the eIPSC amplitudes at the end of the 10 min forskolin application to establish a new baseline and find significant O-GlcNAc iLTD (81.7 ± 9.0% of new baseline (b-c comparison), p=0.005, paired t-test).

To determine if there is any effect of forskolin or GlcN+TMG on presynaptic release probability, we analyzed the paired-pulse ratio (PPR), an indirect measure of presynaptic release probability, during baseline, GlcN+TMG and forskolin. No significant differences were detected in CA1 (Fig. 4Aiv, p=0.69, RM-ANOVA) or in the dentate gyrus (Fig. 4Biv, p=0.46, RM-ANOVA), indicating that a presynaptic mechanism is not involved.

Next, we performed the experiment in reverse order, increasing O-GlcNAc with GlcN+TMG prior to driving phosphorylation with forskolin. We recorded from both from CA1 pyramidal cells and DGCs and applied GlcN+TMG for 10 min followed by forskolin for 10 min. Similar to above, eIPSC amplitudes during forskolin and GlcN+TMG were normalized to baseline and compared (Fig. 5Ai, p<0.0001, RM-ANOVA; 5Bi; p=0.009 RM-ANOVA). A 10 min exposure to GlcN+TMG induced O-GlcNAc iLTD in CA1 pyramidal cells (Fig. 5Aii: 65.4 ± 5.2% of baseline transmission, p=0.0002; Šídák’s post hoc test) and in dentate granule cells (Fig. 5Bi, Bii: 82.3 ± 2.6% of baseline transmission, p=0.002, Šídák’s post hoc test). Surprisingly, subsequent application of forskolin reversed the O-GlcNAc iLTD and elicited a potentiation of the eIPSC amplitude in recordings from both CA1 pyramidal cells and DGCs (Figs. 5Ai-iii and 5Bi-iii). To analyze the magnitude of the forskolin-induced eIPSC potentiation, we re-normalized eIPSC amplitudes at the end of the 10 min GlcN+TMG application, establishing a new baseline, and then normalized forskolin values to the new baseline. We found a significant potentiation in CA1 (138.5 ± 7.8% of new baseline (b-c comparison), p= 0.006, paired t-test) and in dentate (143.4 ± 8.8% of new baseline (b-c comparison), p= 0.003, paired t-test) that reverses O-GlcNAc iLTD and in dentate, the potentiation overshoots the original baseline. Furthermore, these results suggest that a prior increase in O-GlcNAc unmasks a possible PKA dependent potentiation of synaptic inhibition that is absent under control conditions.

**Figure 5.**
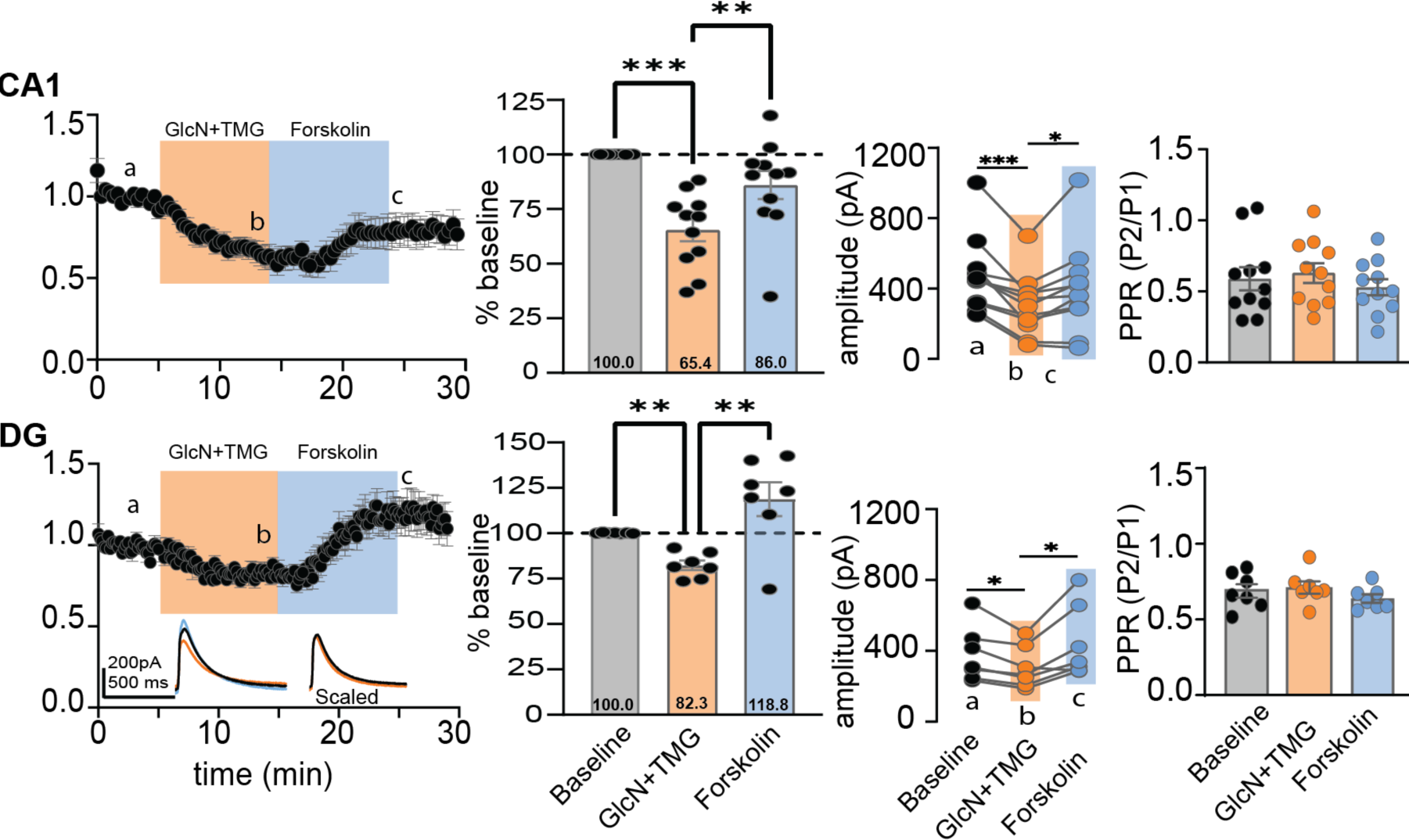
Forskolin reverses O-GlcNAc iLTD in CA1 and dentate. CA1: **Ai.** Normalized data showing average eIPSC amplitude over time during baseline (5 min), bath application of GlcN+TMG (10 min, orange), and forskolin (10 min, blue). *Inset:* overlaid averaged representative eIPSC traces during baseline (black), GlcN + TMG (orange) and forskolin (blue) (left) and scaled (right). **Aii.** Bar chart illustrates the average affect of GlcN+TMG and forskolin on eIPSC amplitude from the original baseline (dotted line): 65.4 ± 5.2%, p=0.0002 (GlcN+TMG) and 86.0% ± 6.3%, p=0.1561 (forskolin) of baseline and each other (p=0.0011); (p<0.0001, RM-ANOVA). **Aiii.** For each individual experiment in Ai, line graphs show the average eIPSC amplitudes during the last 5 min of each condition corresponding to a,b,c in Ai (p < 0.0001, RM-ANOVA, n = 11 cells, 10 rats), Šídák’s: a-b (p=0.0007) and b-c (p=0.028), Baseline: 471.2 ± 64.6 pA, GlcN+ TMG: 304.6 ± 52.0 pA, Forskolin: 397.6± 77.2 pA (mean ± SEM). **Aiv.** Paired-Pulse Ratio between baseline, GlcN+TMG and forskolin showed no significant difference (p=0.06, RM-ANOVA). Dentate: **Bi.** Normalized data showing average eIPSC amplitude over time during baselinebath application of GlcN +TMG (10 min, orange) and forskolin (10 min, blue). *Inset*: overlaid averaged representative eIPSC traces during baseline(black), GlcN + TMG (orange), and forskolin (blue) (left) and scaled (right). **Bii.** Bar chart illustrates the average affect of GlcN+TMG and forskolin on eIPSC amplitude from the original baseline (dotted line): 82.3% ± 2.6%, p=0.002 (GlcN+TMG) and 118.8% ± 9.3%, p=0.2500 (forskolin) and each other (p=0.008); (p=0.009, RM-ANOVA). **Biii.** For each individual experiment in Bi, line graphs show the averaged eIPSC amplitudes for last 5 min of each condition corresponding to a,b,c in **Bi**. (p=0.015, RM-ANOVA, n = 7 cells, 6 rats), Šídák’s: a-b (p=0.0340) and b-c (p=0.038), Baseline: 376.3 ± 58.7 pA, GlcN+ TMG: 306.5 ± 44.2 pA, Forskolin: 443.0± 77.1 pA (mean ± SEM). **Biv.** Paired-Pulse Ratio between baseline, GlcN+TMG and forskolin showed no significant difference (p=0.24, RM-ANOVA). Gray horizontal bars represent the mean ± SEM. Šídák’s post hoc test depicted with asterisks.

### The forskolin dependent increase in eIPSC amplitude is not PKA dependent

To confirm that the forskolin induced potentiation involves PKA dependent phosphorylation (Fig. 6A), we focused our experiments only on CA1 pyramidal cells. We first asked if the PKA inhibitor KT5720 (3 μM) applied for 10 mins before and during forskolin application was able to prevent the forskolin-induced potentiation that reverses the O-GlcNAc iLTD. Experiments with and without KT5720 were interleaved. eIPSC amplitudes during GlcN+TMG and forskolin were normalized to baseline (Fig. 6Bi-Biii) and statistically compared. GlcN+TMG induced O-GlcNAc iLTD and subsequent application of forskolin reversed the O-GlcNAc iLTD and elicited a potentiation of the eIPSC amplitude with (Fig. 6Bii, p=0.004, RM-ANOVA) and without (Fig. 6Biii, p=0.008, RM-ANOVA) KT5720. In the dataset containing KT5720, post hoc Šídák’s multiple comparisons test showed a significant difference between baseline vs. GlcN+ TMG (Fig. 6Bii, p=0.002) and between GlcN+TMG vs. forskolin (Fig. 6Bii, p=0.011). In the dataset without KT5720, post hoc Šídák’s multiple comparisons test showed a significant difference between baseline vs. GlcN+ TMG (Fig. 6Biii, p=0.0011) and between GlcN+TMG vs. forskolin (Fig. 6Biii, p=0.013). To measure the magnitude of the forskolin-induced potentiation, we re-normalized the eIPSC amplitudes at the end of the 10 min GlcN+TMG application to establish a new baseline, and found significant potentiation with (139.7 ± 9.4%, p= 0.004, paired t-test) and without (169.9 ±18.3, p= 0.005, paired t-test) KT5720, and no significant difference between groups (Fig. 6Bi, p= 0.18, unpaired t-test).

**Figure 6.**
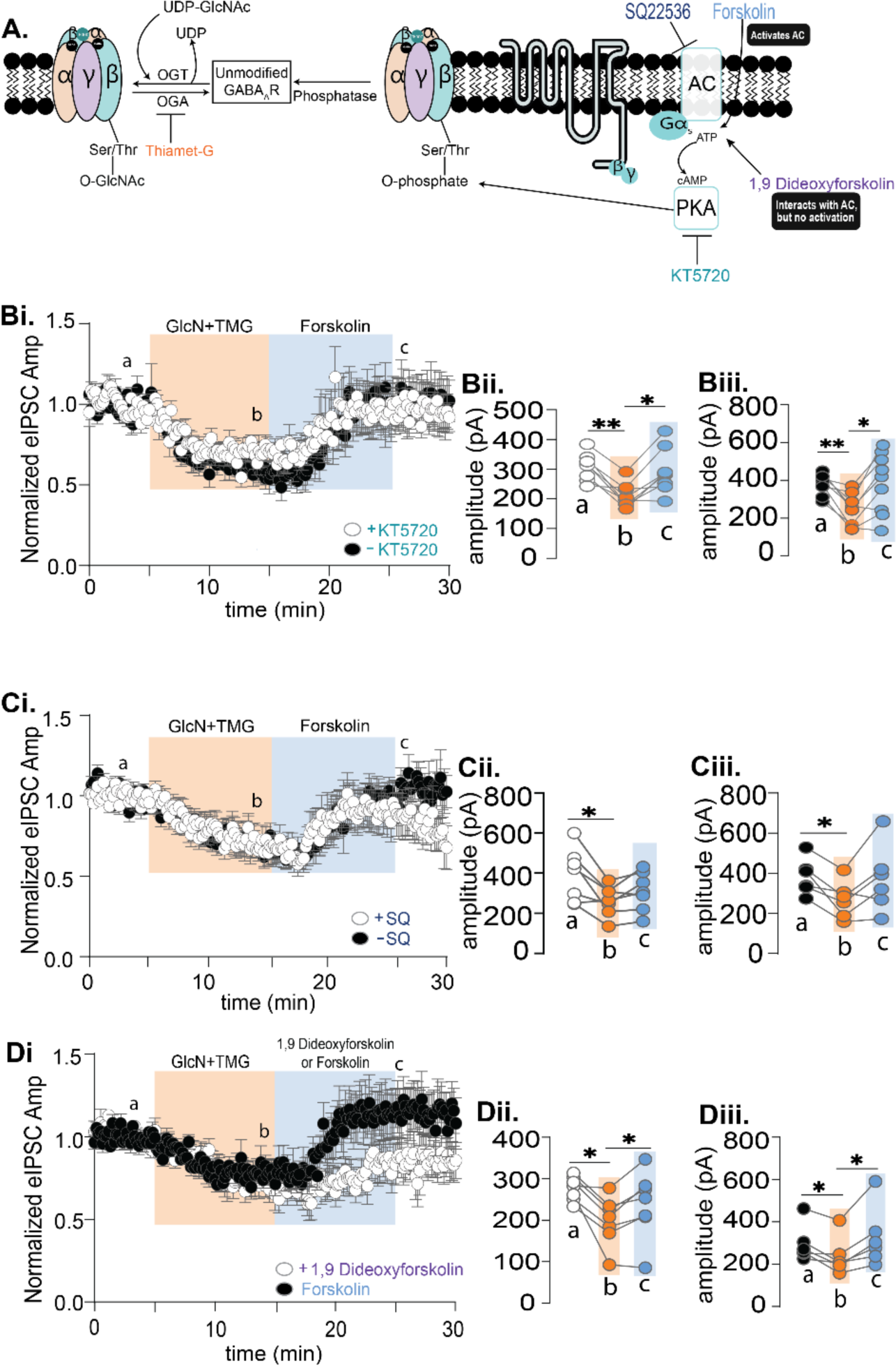
Forskolin induced potentiation of the eIPSC amplitude following an increase in O-GlcNAc is resistant to inhibitors of adenylate cyclase and PKA, suggesting serine phosphorylation is not the mechanism. **A.** Schematic illustrating possible interaction between serine O-GlcNAcylation and PKA-dependent serine phosphorylation of the GABA_A_ receptor with pharmacological activators and inhibitors. **Bi**. Normalized data showing average (±SEM) eIPSC amplitude over time from CA1 pyramidal cells with and without the PKA inhibitor KT5720 (3 μM) during baseline (5 min), GlcN+TMG (10 min, orange) and forskolin application (10 min, blue). KT5720 was applied for 10 mins before and during forskolin. **Bii**,**Biii**. For each individual experiment, line graphs show average eIPSC amplitudes for the last 5 min of each condition corresponding to a,b,c with KT5720 (white) (p=0.004, RM-ANOVA, n= 8 cells, 6 rats), Šídák’s: a-b (p=0.002) and b-c (p=0.011) and without KT5720 (black) (p=0.007, RM-ANOVA, n= 9 cells, 6 rats), Šídák’s: a-b (p=0.0011) and b-c (p=0.0126). **Ci.** Normalized data showing average (±SEM) eIPSC amplitude over time with and without the adenylate cyclase inhibitor, SQ22536 (100 μM) during baseline, GlcN+TMG (10 min, orange) and forskolin application (10 min, blue) **Cii, Ciii** For each individual experiment, line graphs show average eIPSC amplitudes for the last 5 min of each condition corresponding to a,b,c with SQ22536 (white) (p=0.031, RM-ANOVA, n= 7 cells, 6 rats), Šídák’s: a-b (p=0.031) and without SQ22536 (black) (p=0.03, RM-ANOVA, n= 6 cells, 5 rats), Šídák’s: a-b (p=0.020). **Di.** Normalized data showing average (±SEM) eIPSC amplitude over time during baseline, GlcN+TMG (10 min, orange) and application of forskolin or adenylate cyclase inactive analog 1,9 Dideoxyforskolin (10 min, blue). **Dii.** The average amplitudes for last 5 minutes of each condition corresponding to a,b,c with 1,9 Dideoxyforskolin (p= 0.028 RM-ANOVA, n=7 cells, 5 rats), Šídák’s: a-b (p=0.011) and b-c (p=0.037). **Diii.** The average amplitudes for last 5 minutes of each condition with forskolin (p=0.037, RM-ANOVA, n= 6 cells, 4 rats), Šídák’s: a-b (p=0.041) and b-c (p=0.035). Gray horizontal bars represent the mean ± SEM. Šídák’s post-hoc depicted with asterisks.

Since we were unable to block the forskolin mediated potentiation via PKA inhibition, we next targeted adenylate cyclase using the inhibitor, SQ22536 (100 μM). SQ2253 was bath applied for 10 mins before and during forskolin application, and experiments with and without SQ2253 were interleaved. eIPSC amplitudes during GlcN+TMG and forskolin amplitude were normalized to baseline (Fig. 6Ci) and statistically compared (Fig. 6Ci-Ciii). GlcN+TMG induced O-GlcNAc iLTD and subsequent application of forskolin reversed the O-GlcNAc iLTD and elicited a potentiation of the eIPSC amplitude with (Fig. 6Cii, p=0.031, RM ANOVA) and without SQ22536 (Fig. 6Ciii, p=0.03, RM ANOVA). In the dataset with SQ22536, post hoc Šídák’s multiple comparisons test showed a significant difference between baseline vs. GlcN+ TMG (Fig. 6Cii, p=0.031). In the dataset without SQ22536, Šídák’s multiple comparisons test showed a significant difference between baseline vs. GlcN+TMG (Fig. 6Ciii, p=0.020). To measure the magnitude of the forskolin-induced potentiation, we re-normalized the eIPSC amplitudes at the end of the 10 min GlcN+TMG application to establish a new baseline, and found significant potentiation with (122.7± 7.1%, p=0.017, paired t-test) and without (134.8 ± 13.3%, p=0.047, paired t-test) SQ22536, but similar to CA1, there was no significant difference between groups (Fig. 6Ci, p=0.42, unpaired t-test). Being that neither the adenylate cyclase nor PKA inhibitor prevented the forskolin dependent increase in eIPSC following a prior increase in O-GlcNAc, we concluded that this potentiation occurs through another mechanism.

### The inactive adenylate cyclase activator, 1,9 dideoxyforskolin, partially mimics forskolin

While forskolin is known to activate adenylate cyclase ^32–34^, it has many other ‘off-target’ effects ^32,35^ that could be mediating the eIPSC potentiation following O-GlcNAc iLTD. Therefore, we asked whether the adenylate cyclase inactive forskolin analog, 1,9-dideoxyforskolin (50 μM) would mimic the effect of forskolin. Experiments with 1,9-dideoxyforskolin were interleaved with forskolin. eIPSC amplitudes during GlcN+TMG and subsequent application of 1,9 dideoxyforskolin or forskolin were normalized to baseline and statistically compared. GlcN+TMG induced O-GlcNAc iLTD and subsequent application of 1,9 dideoxyforskolin (Fig. 6Dii, p= 0.028, RM-ANOVA) and forskolin (Fig. 6Diii, forskolin p=0.037, RM-ANOVA) reversed the O-GlcNAc iLTD and elicited a potentiation of the eIPSC amplitude. In the 1,9 dideoxyforskolin dataset, Šídák’s multiple comparisons test showed a significant difference between baseline vs. GlcN+ TMG (Fig. 6Dii, p= 0.011), indicating O-GlcNAc iLTD, followed by a significant potentiation upon 1,9 dideoxyforksolin exposure (Fig. 6Dii, p=0.037). In the forskolin dataset, Šídák’s multiple comparisons test showed after O-GlcNAc iLTD (Fig. 6Diii, P=0.041), forskolin exposure induced a significant potentiation (Fig. 6Diii, p=0.035). When comparing the magnitude of the potentiation induced by 1,9 dideoxyforskolin vs forskolin, eIPSC amplitudes were re-normalized at the end of the GlcN+TMG, application to re-establish a new baseline, and found significant potentiation with 1,9 dideoxyforskolin (121.3± 6.3%, p=0.0115, paired t-test) and forskolin (146.5 ± 11.8%, p=0.0075, paired t-test) and no significant difference between groups (Fig. 6Di, p= 0.029, unpaired t-test).

### 5α-pregnane-3α,21-diol-20-one (THDOC) and progesterone reverse the O-GlcNAc-mediated depression of evoked IPSCs, mimicking forskolin

The inability to prevent the forskolin-induced eIPSC potentiation following O-GlcNAc iLTD with adenylate cyclase and PKA inhibitors, and the partial mimic of forskolin’s effect by the inactive adenylate cyclase analog 1,9 dideoxyforksolin, was very puzzling. In searching for a possible explanation, we were intrigued by a report where both forskolin and 1,9-dideoxyforskolin accelerated desensitization of GABA_A_R currents in recordings from amacrine-like cells in carp (Carassius auratus) retina that was resistant to PKA inhibition ^24^. Surprisingly, the neurosteroid, 5α-pregnane-3α,21-diol-20-one (THDOC), which is a structural analog to forskolin, also accelerated GABA_A_R desensitization, mimicking the effect of forskolin ^24^. Further experiments led to the conclusion that forskolin is acting at an allosteric neurosteroid site on GABA_A_Rs. Because GABA_A_Rs in mammalian hippocampus are potently modulated by neurosteroids containing specific subunit combinations ^9^, we sought to determine if the hippocampal neurosteroid, THDOC, also mimics the forskolin-induced eIPSC potentation following O-GlcNAc iLTD.

To test this, we recorded eIPSC from CA1 pyramidal cells and bath applied THDOC (10 μM) before or after GlcN+TMG. The eIPSC amplitudes during THDOC and GlcN+TMG were normalized to baseline and statistically compared (Fig. 7Ai-iii, p=0.003 RM-ANOVA). THDOC application (10 min) led to a slight but not significant depression in eIPSC amplitude (Fig. 7Aii: 83.4 ± 6.9% of baseline transmission, p=0.153) similar to the lack of significant effect of forskolin on baseline transmission (see Fig. 4Ai, Bi). Subsequent application of GlcN+TMG induced O-GlcNAc iLTD (Fig. 7Aii: 56.9 ± 10.3% of baseline transmission, p=0.016).

**Figure 7.**
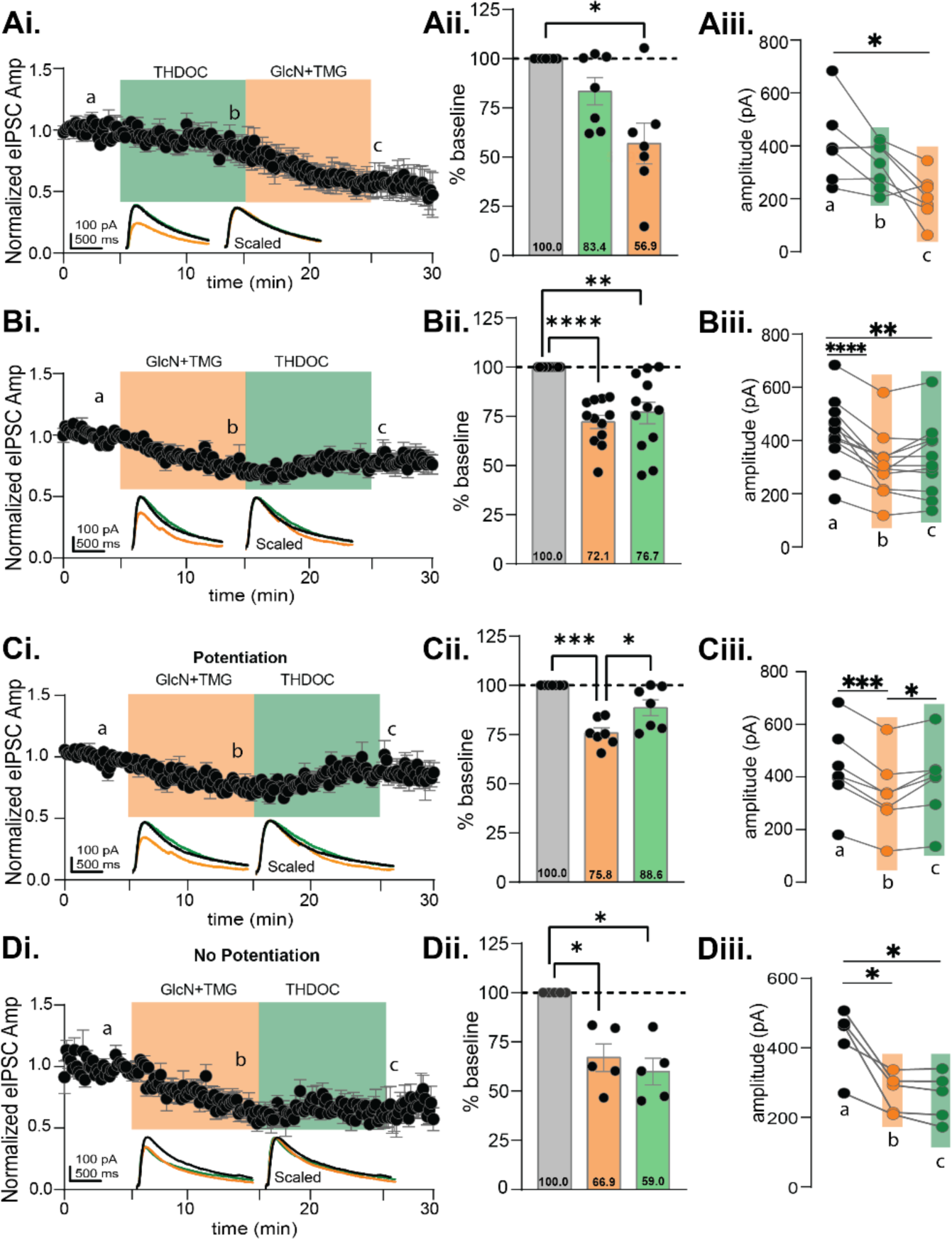
5α-pregnane-3α,21-diol-20-one (THDOC) reverses the O-GlcNAc iLTD, mimicking forskolin in CA1 pyramidal cells. **Ai.** Normalized data showing average eIPSC amplitude during THDOC (10 min, green) followed by GlcN+TMG (10 min, orange). **Aii.** Average THDOC and GlcN+TMG eIPSC amplitude from the original baseline (gray bar, dotted line): 83.4 ± 6.9%, p=0.153 (THDOC) and 56.9%± 10.3%, p=0.016 (GlcN+TMG) and each other (p=0.093); (p=0.0032, RM-ANOVA). **Aiii.** The average eIPSC amplitudes during the last 5 min of each condition corresponding to a,b,c in Ai (p= 0.0059, RM-ANOVA, n = 7 cells, 7 rats), Šídák’s: a-c (p=0.032). **Bi.** Normalized data showing average eIPSC amplitude during GlcN+TMG (10 min, orange) followed by THDOC (10 min, green). **Bii.** Average GlcN+TMG and THDOC eIPSC amplitude from the original baseline (gray bar, dotted line): 72.1% ± 3.3% (GlcN+TMG), p< 0.0001 and 76.7% ± 5.5% (THDOC), p=0.003 and each other (p=0.617); (p<0.0001, RM-ANOVA). **Biii.** The average eIPSC amplitudes during the last 5 min of each condition corresponding to a,b,c in Bi (p <0.0001, RM-ANOVA, n = 12 cells, 12 rats), Šídák’s: a-b (p<0.0001), a-(p=0.005). *Potentiated:* **Ci.** Normalized data showing average eIPSC amplitude during GlcN+TMG followed by THDOC. **Cii.** Average GlcN+TMG and THDOC eIPSC amplitude from the original baseline (gray bar, dotted line): 75.8% ± 2.6% (GlcN+TMG), p= 0.0002 and 88.6% ± 4.0% (THDOC), p=0.089 and each other (p=0.034); (p=0.0002, RM-ANOVA). **Ciii.** Last 5 min average amplitudes corresponding to a,b,c (p=0.0003, RM-ANOVA, n = 7 cells, 7 rats), Šídák’s: a-b (p=0.0002), b-c (p=0.036). *Non-potentiated:* **Di.** Normalized data showing average eIPSC amplitude during GlcN+TMG followed by THDOC. **Dii.** Average GlcN+TMG and THDOC eIPSC amplitude from the original baseline (gray bar, dotted line): 66.9% ± 6.9% (GlcN+TMG), p= 0.026 and 59.9% ± 6.8% (THDOC), p=0.012 and each other (p=0.422); (p=0.0014, RM-ANOVA). **Diii.** Last 5 min average amplitudes corresponding to a,b,c (p=0.0099, RM-ANOVA, n = 5 cells, 5 rats), Šídák’s: a-b (p=0.041), a-c (p=0.025). *Inset:* overlaid averaged representative eIPSC traces during baseline (black), GlcN + TMG (orange), THDOC (green) (left) and scaled (right). Gray horizontal bars represent mean ± SEM. Šídák’s post-hoc depicted with asterisks.

Next, we performed the experiment in the reverse order to specifically ask whether THDOC mimics forskolin by eliciting a potentiation of the eIPSC amplitude when applied after GlcN+TMG (Fig.7Bi). The eIPSC amplitudes recorded from CA1 pyramidal cells during GlcN+TMG and THDOC were normalized to baseline and statistically compared (Fig. 7Bi-iii, p<0.0001, RM-ANOVA). Bath application of GlcN+TMG induced O-GlcNAc iLTD (Fig. 7Bii: 72.1 ± 3.3% of baseline transmission, p<0.0001), but subsequent application of THDOC yields no additional change in eIPSC amplitude (Fig. 7Bii: 76.7 ± 5.5% of baseline transmission, p=0.003), and is not different from eIPSC amplitude following GlcN+TMG (p=0.2738, paired t-test). There was a significant difference between baseline vs. GlcN+ TMG (Fig. 7Biii, p<0.0001) and baseline vs. THDOC (Fig. 7Biii, p=0.005).

However, similar to the cell-to-cell variability we observed with forskolin (e.g., Fig. 4Aii), we noted variability in the response to THDOC (Fig. 7Bii), with eIPSCs recorded from some cells having a clear potentiation (Fig. 7Ci-iii, n=7/12), while others had no change (Fig.7 Di-iii, n=5/12 cells). To measure the effect of THDOC in these two populations, we re-normalized eIPSC amplitudes at the end of the 10 min GlcN+TMG application to establish a new baseline, then THDOC values were normalized to the new baseline. Similar to forskolin (Fig.4Ai-iii), we found a significant potentiation of the eIPSC amplitude in this subset of cells (116.4 ± 5.3%, p= 0.021, paired t-test, n=7/12). In the remaining cells, (Fig. 7Di-iii), we found no further change in eIPSC amplitude (90.0 ± 5.5%, p= 0.15, paired t-test) beyond what occurred following GlcN+TMG application.

To firm up the above results, we felt it important to test a second steroid, progesterone, known to modulate GABA_A_Rs, to see if the potentiation following GlcN+TMG is again mimicked. The eIPSC amplitudes recorded from CA1 pyramidal cells during progesterone and GlcN+TMG were normalized to baseline and statistically compared (Fig. 8Ai-iii, p=0.047, RM-ANOVA). Similar to forskolin and THDOC, progesterone exposure induced no significant change from baseline (Fig. 8Aii: 92.2 ± 7.3% of baseline, p=0.701) and subsequent GlcN+TMG induced O-GlcNAc iLTD (Fig. 8Aii: 76.9% ± 9.1% of baseline, p=0.126). A significant difference also existed between progesterone vs. GlcN+TMG (Fig. 8Aiii, p=0.008). When performing the experiment in reverse, the eIPSC amplitudes recorded from CA1 pyramidal cells during GlcN+TMG and progesterone were normalized to baseline and statistically compared (Fig 8Bi, p<0.0001, RM-ANOVA). GlcN+TMG induced O-GlcNAc iLTD (Fig. 8Bii: 68.7% ± 3.8% of baseline transmission, p<0.001,) but subsequent application of progesterone yields no additional change in eIPSC amplitude (Fig. 8Bii: 75.3% ± 4.9% of baseline transmission, p=0.0011), and is not different from eIPSC amplitude following GlcN+TMG (p=0.1705, paired t-test). There was a significant difference between baseline vs. GlcN+ TMG (Fig. 8Biii, p=0.0004).

**Figure 8.**
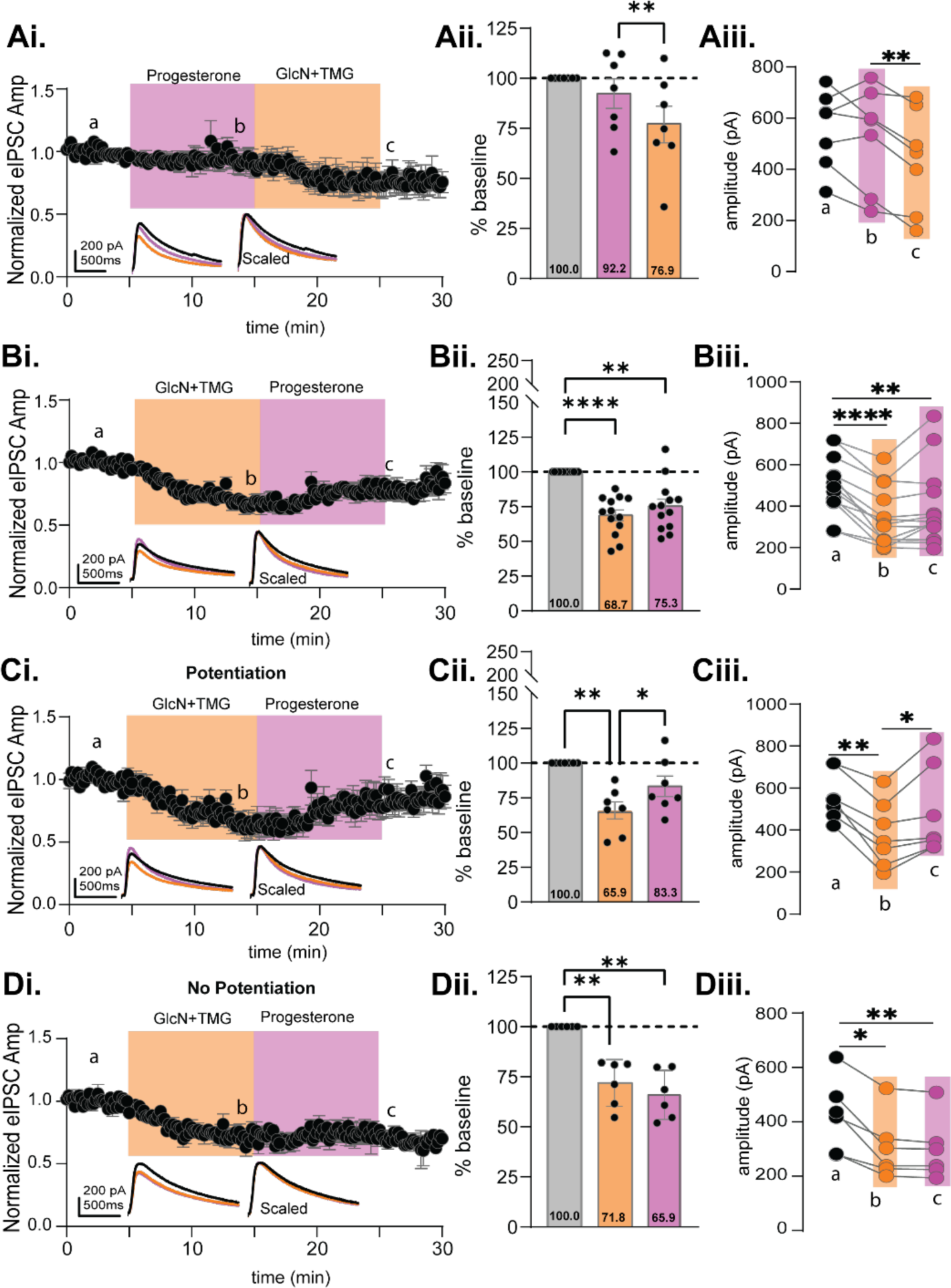
Progesterone also reverses O-GlcNAc iLTD, mimicking forskolin in CA1 pyramidal cells. **Ai.** Normalized data showing average eIPSC amplitude during progesterone (10 min, pink) followed by GlcN+TMG (10 min, orange). **Aii.** Average progesterone and GlcN+TMG eIPSC amplitude from the original baseline (gray bar, dotted line): 92.2% ± 7.3% (Progesterone), p=0.701 and 76.9% ± 9.1% (GlcN+TMG), p=0.126 (GlcN+TMG) and each other (p=0.008); (p=0.047, RM-ANOVA). **Aiii.** The average eIPSC amplitudes during the last 5 min of each condition corresponding to a,b,c in Ai (p= 0.0328, RM-ANOVA, n = 7 cells, 7 rats). Šídák’s: b-c (p=0.008). **Bi.** Normalized data showing average eIPSC amplitude during GlcN+TMG (10 min, orange) followed by progesterone (10 min, pink). **Bii.** Average GlcN+TMG and progesterone eIPSC amplitude from the original baseline (gray bar, dotted line): 68.7% ± 3.8%, p<0.001 (GlcN+TMG), and 75.3% ± 4.9% (progesterone), p=0.0011 and each other (p=0.428); (p<0.0001, RM-ANOVA). **Biii.** The average eIPSC amplitudes during the last 5 min of each condition corresponding to a,b,c in Bi (p= 0.0007, RM-ANOVA, n= 13 cells, 10 rats) Šídák’s: a-b (p=0.0004). *Potentiated:* **Ci.** Normalized data showing average eIPSC amplitude during GlcN+TMG followed by progesterone. **Cii.** Average GlcN+TMG and progesterone eIPSC amplitude from the original baseline (gray bar, dotted line): 65.9% ± 6.2% (GlcN+TMG), p= 0.0045 and 83.3% ± 7.1% (progesterone), p=0.175 and each other (p=0.019); (p=0.002 RM-ANOVA). **Ciii.** Last 5 min average amplitudes corresponding to a,b,c (p=0.002, RM-ANOVA, n = 7 cells, 7 rats) Šídák’s: a-b (p=0.0019), b-c (p=0.037). *Non-potentiated:* **Di.** Normalized data showing average eIPSC amplitude during GlcN+TMG followed by progesterone. **Dii.** Average GlcN+TMG and progesterone on eIPSC amplitude from the original baseline (gray bar, dotted line): 71.8% ± 4.7% (GlcN+TMG), p= 0.006 and 65.9% ± 5.0% (progesterone), p=0.0031 and each other (p=0.591); (p<0.0001, RM-ANOVA). **Diii.** Last 5 min average amplitudes corresponding to a,b,c (p=0.004, RM-ANOVA, n = 6 cells, 6 rats), Šídák’s: a-b (p=0.015), a-c (p=0.0096). *Inset:* overlaid averaged representative eIPSC traces during baseline (black), GlcN + TMG (orange), progesterone (pink) (left) and scaled (right). Gray horizontal bars represent mean ± SEM. Šídák’s post hoc depicted with asterisks.

Similar to THDOC, we noted variability in the progesterone response (Fig. 8Bii), eIPSCs recorded from some cells having a potentiation (Fig. 8Ci-Ciii, n = 7/13), and others, no potentiation (Fig. 8Di-Diii, n =6/13). To measure the effect of progesterone in these two populations, we re-normalized eIPSC amplitudes at the end of the 10 min GlcN+TMG application to establish a new baseline, then progesterone values were normalized to the new baseline. Similar to forskolin (Fig.4Ai-iii) and THDOC (Fig.7Ci-iii), we found a significant potentiation of the eIPSC amplitude in this subset of cells (134.4 ± 7.6%, p= 0.0041, paired t-test, n=7/13). In the remaining cells, (Fig. 8Di-iii), we found a slight but significant decrease (96.4 ± 1.3%, p=0.0443, paired t-test).

## Discussion

Understanding the mechanisms that modulate the strength of inhibitory transmission at GABAergic synapses is essential to understanding the excitation/inhibition balance critical for brain function. We previously reported that pharmacologically increasing the post-translational modification O-GlcNAcylation rapidly depresses spontaneous IPSC frequency and amplitude, and the amplitude of miniature IPSCs, suggesting the mechanism underlying the synaptic depression is postsynaptic^22^. We also reported a long-lasting depression of electrically evoked IPSC amplitude^22^, representing a novel form of LTD of synaptic inhibition and referred to here as O-GlcNAc iLTD. In the current study, we extend these initial findings by investigating whether increasing O-GlcNAcylation triggers GABA_A_R endocytosis during expression of O-GlcNAc iLTD and how O-GlcNAcylation and serine phosphorylation might interact in the modulation of GABAergic inhibition on principle cells in hippocampus.

A growing list of mechanisms regulate GABA_A_R membrane stability and trafficking during long-term changes in strength of inhibitory transmission^36–39^. Under some conditions, GABA_A_R receptors can undergo clathrin/dynamin-mediated endocytosis involving PKA-dependent phosphorylation of serine 408 (S408) and S409 on β1 and β3 subunits^40–43^. Similarly, depending upon the subunit composition, PKC dependent serine phosphorylation also modulates endocytosis, leading to decreased plasma membrane GABA_A_R density^44–47^. Because O-GlcNAcylation also occurs on serine residues, and O-GlcNAc LTD of excitatory transmission likely involves endocytosis of AMPARs ^20,27^, it seemed probable that GABA_A_R endocytosis occurs during expression of O-GlcNAc iLTD. Surprisingly, the actin stabilizer, jasplakinolide did not prevent expression of O-GlcNAc iLTD, but pharmacologically inhibiting dynamin-dependent trafficking with dynasore prevented O-GlcNAc iLTD expression in about 50% of recorded cells. In those specific cells, a significant potentiation of the eIPSC amplitude was observed which is reminiscent of previous results where dynamin inhibition in cultured hippocampal neurons resulted in accumulation of postsynaptic GABA_A_Rs and increased mIPSC amplitude^42^. The variable effect of dynasore on O-GlcNAc iLTD suggests that the expression mechanism is complex. This is not too surprising since the specific combination of scaffolding proteins that interact with GABA_A_Rs varies among inhibitory synapses in specific brain regions, neuron types, and even within specific regions within the same neuron^37^. Therefore, it is possible that the precise mechanism underlying O-GlcNAc iLTD may also be variable. Like AMPARs, GABA_A_Rs are highly dynamic within inhibitory synapses and can rapidly undergo lateral diffusion, causing depression of inhibitory transmission^48–51^. Whether increasing O-GlcNAcylation alters GABA_A_R subunit interaction with gephyrin or other scaffolding proteins to stimulate lateral diffusion that underlies expression of O-GlcNAc iLTD is currently unknown and is an area of needed future investigation.

The known interplay, and sometimes competition, between serine O-GlcNAcylation and phosphorylation on key proteins ^15,52–54^ led us to further explore how a possible interaction might impact the strength of inhibitory transmission. A notable example of this O-GlcNAcylation-phosphorylation interaction is competition for the same serines on Tau where increasing O-GlcNAc prevents hyperphosphorylation of Tau and development of tangles in Alzheimer’s disease^53^. Serine phosphorylation has complex effects on GABA_A_R function, including impacting how channel function is modulated by benzodiazepines, barbiturates, and neurosteriods^8,10,12^. An interaction between O-GlcNAcylation and phosphorylation would add to the complexity, and we specifically focused on PKA dependent phosphorylation using the adenylate cyclase activator, forskolin. As mentioned previously, this strategy has been used to investigate PKA dependent potentiation of excitatory transmission in hippocampus^29–31^. While forskolin had a variable effect on the amplitude of the eIPSC during baseline transmission, causing depression in some cells and potentiation in others with no statistically significant overall effect, it did not impact the magnitude of subsequently induced O-GlcNAc iLTD. This is consistent with no interaction between a prior increase in PKA dependent phosphorylation with subsequent O-GlcNAc modification. However, forskolin unexpectedly reversed the polarity of eIPSC amplitude when applied during expression of O-GlcNAc iLTD, eliciting a significant potentiation of the eIPSC amplitude that in some cells even overshot the original baseline. It is important to note that this unexpected potentiation of the eIPSC amplitude occurs at inhibitory synapses in both CA1 and dentate gyrus, suggesting a general mechanism that may not be too dependent on a specific subunit composition. Moreover, this highly interesting finding suggests the possibility that direct O-GlcNAc modification of GABA_A_R subunits, and/or specific scaffolding proteins, changes the GABA_A_R confirmation in a way that unmasks this potentiating effect of forskolin. A further surprise was that this forskolin-dependent eIPSC potentiation was not prevented by pharmacological inhibition of either adenylate cyclase nor PKA, and was mimicked by the adenylate cyclase inactive forskolin analog 1,9 dideoxyforskolin. Thus, this interaction of O-GlcNAcylation and forskolin is not a consequence of PKA mediated phosphorylation.

Clearly, forskolin is working through some other mechanism to potentiate inhibitory transmission following a prior increase in O-GlcNAcylation. Ironically, a previous report demonstrated forskolin-dependent increase in GABA_A_R desensitization in carp retina through a non-PKA dependent mechanism that involved acting directly on a neurosteroids site ^24^. Neurosteroids and metabolites of progesterone positively modulate GABA_A_Rs in a dose dependent manner by acting on synaptic and extrasynaptic GABA_A_Rs, and their mechanism of action can be enhanced or diminished depending on the activation or inhibition of serine phosphorylation of GABAARs as well as GABA_A_R subunit expression^10,12,55^. Furthermore, neurosteroids, such as THDOC, occupy a binding pocket in the transmembrane region that can involve conserved threonines in the α1 and α5 subunits. Importantly, threonines can also undergo O-GlcNAc modification similar to serines.

Our data showing that both THDOC and progesterone reversed O-GlcNAc iLTD in recordings from CA1 pyramidal cells, while having no significant effect on baseline eIPSC amplitude, precisely mimics the effect of forskolin. These findings supports the interpretation that forskolin is acting at the neurosteroid site on synaptic GABA_A_Rs. Perhaps most exciting is the observation that these steroids induced no significant effect on baseline eIPSC amplitude, but could only modulate the strength of synaptic inhibition following a prior increase in O-GlcNAcylation. Thus, O-GlcNAc modification enables synaptic GABA_A_Rs to be modulated by neurosteroids and potentiate the eIPSC amplitude thereby reversing the polarity of the iLTD. It is important to point out that not all recorded cells exhibit this reversal in polarity of the eIPSC amplitude, suggesting that there may be a GABA_A_R subunit confirmation preference and/or subunit combination preference, a concept supported by the varying subunit composition of GABA_A_Rs across the same cell types and across different locations on the same cell, leading to different responses upon exposure to allosteric modulators^56^. In addition, our findings indicate that somehow O-GlcNAcylation enhances access to the neurosteroid site on GABA_A_Rs for both forskolin and allosteric modulators to act. Because GABA_A_Rs are therapeutic targets for drugs used in the treatment of neurological and neuropsychiatric conditions, understanding how they are modulated by O-GlcNAcylation has clinical implications, particularly if O-GlcNAc interferes with or enhances their efficacy. Future work is needed to determine whether GABA_A_R subunits and/or scaffolding proteins are directly O-GlcNAc modified, and if so, which specific serines and/or threonines are modified. Furthermore, understanding how elevated O-GlcNAc impacts synaptic stability and lateral diffusion within the membrane, and how the neurosteroid site becomes more accessible at synaptic receptors.

Collectively, these current results, together with our previously published results^20–22^ not only solidifies O-GlcNAcylation as a critical regulator of both synaptic inhibition and excitation, but also provides highly novel information that O-GlcNAc dictates the polarity of the change in GABA_A_R synaptic current amplitude mediated by endogenous neurosteroids THDOC and progesterone, highlighting O-GlcNAcylation’s ability to modify the effectiveness of allosteric modulators on GABAergic transmission. Thus, our current findings uncovers how protein O-GlcNAcylation can possibly serve as a gauge for the potency of synaptic inhibition and its modulation by allosteric modulators and novel therapeutic agents.

## Materials and Methods

All experimental procedures were approved by the Medical University of South Carolina Institutional Animal Care and Use Committee and follow the National Institutes of Health experimental guidelines.

### Hippocampal slice preparation

Male and female Sprague Dawley rats (age 3–7 weeks; Charles River Laboratories) were anesthetized with isoflurane, decapitated, and brains removed; 400 μm coronal slices from dorsal hippocampus were prepared on a VT1200S vibratome (Leica Biosystems) in oxygenated (95%O2/5%CO2) ice-cold, high sucrose cutting solution (in mM as follows: 85.0 NaCl, 2.5 KCl, 4.0 MgSO4, 0.5 CaCl2, 1.25 NaPO4, 25.0 glucose, 75.0 sucrose). After cutting, slices were held at room temperature for 1 to 5 hr in a submersion chamber with continuously oxygenated standard ACSF (in mM as follows: 119.0 NaCl, 2.5 KCl, 1.3 MgSO4, 2.5 CaCl2, 1.0 NaH2PO4, 26.0 NaHCO3, 11.0 glucose).

### Electrophysiology

All recordings were performed in a submersion chamber with continuous perfusion of oxygenated standard ACSF. The blind patch technique was used to acquire interleaved whole-cell recordings from CA1 pyramidal neurons and dentate granule cells. Neuronal activity was recorded using an Axopatch 200B amplifier and pClamp10.7 acquisition software (Molecular Devices, Sunnyvale, CA). Signals were filtered at 5 kHz and digitized at 10 kHz (Digidata 1440). Patch pipettes (BF150-110 HP; Sutter Instruments, Novato, CA) were pulled on a Sutter P-97 (Sutter Instruments, Novato, CA) horizontal puller to a resistance between 3-5 MΩ. Spontaneous IPSCs were recorded using CsCl internal solution (in mM: 140.0 CsCl, 10.0 EGTA, 5.0 MgCl2, 2.0 Na-ATP, 0.3 Na-GTP, 10.0 HEPES; E_Cl_ = 0 mV). Evoked GABA_A_R currents were recorded using Cs-gluconate internal solution (in mM: 100.0 Cs-gluconate, 0.6 EGTA, 5.0 MgCl2, 2.0 Na-ATP, 0.3 Na-GTP, 40.0 HEPES; E_Cl_ = −60 mV) with a twisted nichrome wire bipolar electrode positioned in stratum radiatum to activate Schaffer collateral axons (0.1 Hz, 100 μs duration) when recording in CA1 and positioned in the medial performant path when recording in the dentate gyrus. GABA_A_R currents were pharmacologically isolated with bath perfusion of DNQX (10μM; Hello Bio) and R-CPP (5 μM; Hello Bio). Recordings were performed at a 10 mV test pulse at the end of each sweep to monitor series resistance and was excluded if there was more than a 20% change during the experiment. Stability of series resistance was verified using post-hoc scaling of averaged waveforms before and after pharmacologically increasing O-GlcNAcylation and after Forskolin, KT5720, SQ22536, 1,9 Dideoxyforskolin, Jasplakinolide and THDOC exposure.

### Chemicals

Forskolin (Hello Bio) and 1,9 Dideoxyforskolin (Sigma-Aldrich) were prepared as a 50 mM stock in DMSO and stock was added to external solution for a final concentration of 50 μM. KT5720 (Hello Bio) was prepared as a 25 mM stock in DMSO and stock was added to external solution for a final concentration of 3 μM. SQ22536 (Tocris), 5α,21-pregnane-3α,21-diol-20-one (THDOC) (Sigma-Aldrich, mixed and sonicated), Progesterone (Sigma-Aldrich) was prepared as a 100 mM stock in DMSO and the stock was added to external solution for a final concentration of 100 μM, 10 μM, and 1 μM respectively. Jasplakinolide (Hello Bio) was prepared as a 1 mM stock in DMSO and stock was added to Cs-gluconate internal solution for a final concentration of 2 µM. Dynasore (Sigma-Aldrich) was prepared as a 100 mM stock in DMSO and stock was added to external solution for a final concentration of 80 μM. The actin stabilizer jasplakinolide (2μM, Hello Bio)^57,58^ was included in the pipet solution to prevent GABA_A_R endocytosis. PKA dependent serine phosphorylation was triggered by bath application of the adenylate cyclase activator, forskolin (50μM, Hello Bio) ^31^ determine if the O-GlcNAc induced synaptic depression is prevented.

### Alterations in the levels of O-GlcNAcylation

As in our previously published papers^20–22^, the HBP substrate glucosamine (GlcN, 5 mM; Sigma) and OGA inhibitor, Thiamet-G (TMG) (10 min) (1 μM; Chem Molecules) were combined to acutely and robustly increase O-GlcNAcylation via bath application *in vitro*, ensuring a long-lasting effect.

### Statistical analysis

Recordings were analyzed using Clampfit 11.2. GraphPad Prism 10.0.2, La Jolla, CA, was used for all analyses and outliers were excluded. Statistical significance was detected via two-tailed paired or unpaired Student’s t-tests (parametric), or Wilcoxon matched-pairs signed rank test (non-parametric, paired) when comparing two-groups. For percent baseline calculations, the amplitudes of the last five minutes of each drug treatment for each cell were normalized to baseline or GlcN+TMG, as the new baseline, averaged and converted into percentages. Multi-groups were analyzed using RM-ANOVA with post hoc Šídák’s multiple comparisons test (nonparametric). Data are displayed as mean ± SEM and p values considered statistically significant were as follows: p ≤ 0.05, **p ≤ 0.01,***p ≤ 0.001,***p ≤ 0.0001,****p ≤ 0.00001.

## Acknowledgements

This research was supported by NIH NINDS R01AG066489 to LLM and JCC, 1F99NS134163-01 to SMP

## Author Contributions Statement

S.P. and L.L.M. designed the experiments; S.P. conducted experiments; S.P. and L.L.M. analyzed data; S.P, J.C.C., and L.L.M. wrote and revised the manuscript.

## Additional Information

Competing Interests Statement: The authors declare no competing interests.

## Data Availability

The datasets generated and analyzed during the current study are available from the corresponding author on reasonable request.

